# Position- and scale-invariant object-centered spatial selectivity in monkey frontoparietal cortex dynamically adapts to task demand

**DOI:** 10.1101/2022.01.26.477941

**Authors:** Bahareh Taghizadeh, Ole Fortmann, Alexander Gail

**Author notes:** Corresponding author: Alexander Gail. Authors’ contribution: B.T. and A.G. designed and implemented the experiment, and collected the data. B.T. and O.F. analyzed the data. B.T., O.F. and A.G. discussed the results and wrote the paper.

## Abstract

Humans utilize egocentric and allocentric spatial information to guide goal-directed movements. Egocentric encoding is a well-known property of brain areas along the dorsal pathway. We ask if dorsal stream reach planning areas like parietal reach region (PRR) and dorsal premotor cortex (PMd) also encode object-centered (allocentric) information. During two consecutive instructed delay periods, rhesus monkeys first memorized an object-relative target position and then planned a reach to this position after the object re-occurred at variable positions with potentially different size. In both areas, we find predominant object-centered encoding during visual memory, which is invariant to object position and object size, and predominant egocentric encoding during reach planning. Such dynamic transition from allo- to egocentric encoding within individual dorsal stream areas contrasts the idea of task-independent functional segregation between processing pathways. Instead, demand-specific local computations might facilitate spatial cognition in dynamic environments to facilitate motor planning towards objects changing their location.

**Significance statement:** Human and non-human primates interact with their environment by manipulating objects. This requires planning and executing reaches to transport the hand to object-relative positions close to or on the object. Previous studies showed that frontal and parietal lobe areas encode reach goals in coordinate systems anchored to different parts of the subjects’ body, such as hand position or gaze direction, and modulated by pose. We show that neurons in the same areas can encode object-centered allocentric spatial information, independent of object location and object size, or egocentric information, depending on dynamically changing task demands. Such dynamic adaptation seems inconsistent with a functional segregation for ego- and allocentric encoding between processing streams.

## Introduction

Allocentric (subject-independent) spatial cognition is a fundamental skill supporting navigation (Moser et al., 2008), spatial judgment (Aagten-Murphy and Bays, 2019; Galati et al., 2000; Vallar et al., 1999; Zaehle et al., 2007), and goal-directed movement behavior (Byrne et al., 2010). Since primary spatial sensory inputs are egocentric, e.g. determined by visual retinotopy and tactile somatotopy, allocentric encoding must be the result of neural computation. Yet, brain regions, especially the roles of ventral versus dorsal stream processing, and neurocomputational mechanisms of allocentric encoding are still under debate (Chen and Crawford, 2020; Deneve and Pouget, 2003; Filimon, 2015). Neurophysiology data at the single neuron level in the context of allocentric goal-directed reaching is lacking. Here we ask if object-centered encoding can be observed during goal-directed reach planning in frontoparietal areas of the dorsal stream, i.e., areas which are mostly associated with egocentric frames of reference, e.g., (Battaglia-Mayer et al., 2003; Bremner and Andersen, 2014; Chang and Snyder, 2010).

Natural reaches typically are directed towards physical objects. Geometrical features such as shape and size of that object should be incorporated in movement planning to successfully direct the hand towards a suited part of the object. We may pick up a stick at different positions along its length, depending on the intended use. Such object-oriented reach goal may only depend on object properties and hence can be defined independently of the spatial positioning of the object relative to the subject. This would mark a case of reach-associated, yet object-centered allocentric spatial processing for the localization of the target position of the reach. Additionally, the inherently egocentric changes in body-configuration needed to accomplish the reach will become relevant for planning and implementing it. It is an open question if dynamically changing allo- and egocentric spatial cognitive demands for planning goal-directed movements are fulfilled in separate specialized brain regions, or if the same local neural network within an area can reflect both types of encoding.

Human neuroimaging studies on allocentricity in the context of perceptual judgment tasks (with verbal or button-press responses) suggest an anatomical segregation, where the dorsal stream ‘vision-for-action’ processing is predominantly egocentric and the ventral stream ‘perceptual’ and ‘object-recognition’ processing encompasses also allocentric representations (Galati et al., 2000; Neggers et al., 2006). Yet, it may be task dependent which processing predominates and it has been shown that allocentric spatial judgments can activate networks comprising areas of both dorsal and ventral streams (Committeri et al., 2004; Zaehle et al., 2007). Human neuroimaging studies on target localization partly also indicated activation of overlapping regions of the frontoparietal network for planning and guiding goal-directed reach and saccade movements relative to egocentric and allocentric references (Chen et al., 2018, 2014, 2011; Thaler and Goodale, 2011a, 2011a, 2011b). However, the role of dorsal stream in allocentric processing is not fully clear from imaging studies. While allocentric compared to egocentric encoding of reach target distance and direction led to higher blood-oxygen-level-dependent (BOLD) activity in dorsal premotor cortex (PMd) and right posterior intraparietal sulcus of human subjects (Thaler and Goodale, 2011c), others did not find such preference along the dorsal stream (Chen et al., 2018; 2014).

Single neuron recordings in monkeys revealed how sensory information is transformed into motor goal information within the frontoparietal network during reach planning (Cui and Andersen, 2011). Neurons in monkey posterior medial intraparietal cortex (parietal reach region, PRR) and PMd are selective for the spatial location of visual cues indicating the target for a reach, but also cue-independent reach goals (Cisek and Kalaska, 2005; Crammond and Kalaska, 1994; Gail et al., 2009; Gail and Andersen, 2006; Hwang and Andersen, 2012; Riehle and Requin, 1989; Snyder et al., 1997; Westendorff et al., 2010; Wise et al., 1997). In monkeys that aim their reach at visually instructed positions, while controlling gaze independently, different frames of reference have been described, all of which are egocentric. Many studies reported predominant gaze-centered (Batista et al., 2007, 1999; Bremner and Andersen, 2014; Buneo et al., 2002; Chang and Snyder, 2010; Cohen and Andersen, 2002; Marzocchi et al., 2008) or predominant hand-centered (Bremner and Andersen, 2012; Caminiti et al., 2015, 1991, 1990; Johnson et al., 1996) encoding of reach goals. Encoding within these areas is typically not exclusive but rather intermediate with mixed (Chang and Snyder, 2012; McGuire and Sabes, 2011) and more complex selectivities (Pesaran et al., 2006), also found in humans (Zhang et al., 2017).

The prevailing view of dorsal stream processing that emerged from these neurophysiological findings is that different visual and somatosensory inputs, which are of egocentric nature, become integrated in the parietal multisensory association cortex to compute reach-relevant spatial information in different egocentric reference frames. Different anatomical nodes have a predominance for reference frames being centered on different body parts (Batista et al., 1999; Bremner and Andersen, 2012; Chang and Snyder, 2012, 2010), and the result of such feedforward integration (Deneve and Pouget, 2003; Zipser and Andersen, 1988) is supposed to be fed to more motor-related areas in the frontal lobe. Yet, if also allocentric spatial cognitive processing in the context of goal-directed reaching exists in parietal cortex is unclear, since single-cell electrophysiology studies directly comparing egocentric and allocentric reference frames during reach planning do not exist.

Here we directly compare posterior medial intraparietal cortex (parietal reach reagion, PRR) and dorsal premotor cortex (PMd) at the single neuron level in rhesus monkeys in an object-centered allocentric reach task that sequentially mandates spatial target memory and reach planning. Two experiments tested position- and size-invariant object-centered encoding. In contrast to the prevailing view, we report that neurons in PRR and PMd encode cue and reach target in object-centered as well as egocentric reference frame and the predominating reference frame in both areas is dynamically adjusted to the cognitive needs.

## Results

Two male rhesus monkeys (Macaca mulatta) were trained to perform memory guided reaches towards variable positions on an elongated object. The object had variable position relative to the animal, allowing to dissociate object-centered (allocentric) locations from body-centered (egocentric) reach goal locations (**Fig. 1**). Two instructed delays allowed separately investigating spatial frames of reference during visual memory and reach planning, respectively.

**Figure 1:**
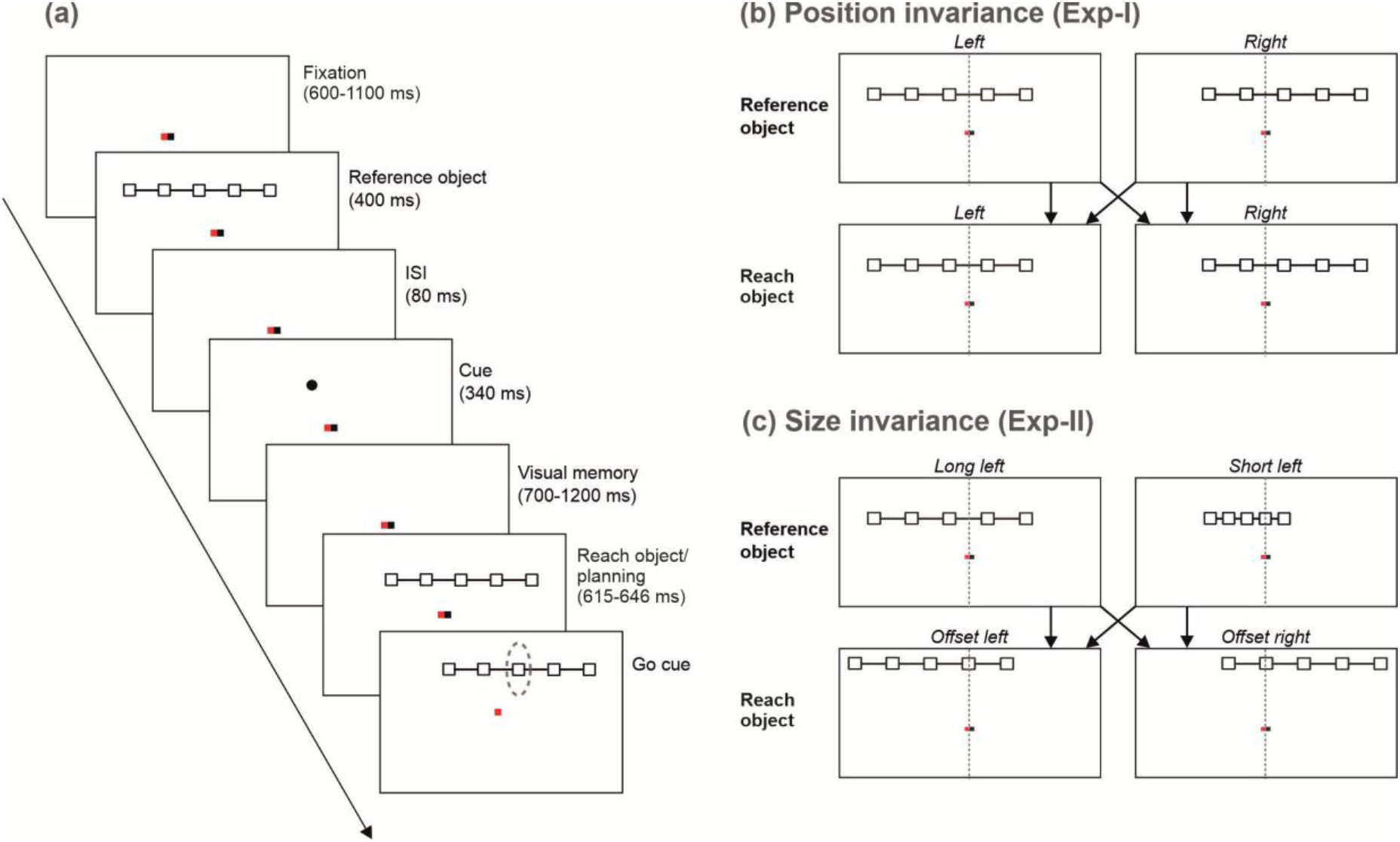
Object-based reach planning task. **(a) Time course**. After acquiring and holding successful ocular fixation of the central red square and touching of the white square during baseline, an array of five interconnected boxes (reference object) is presented randomly to the left or right of the screen center. The reference object is followed by a brief cue presentation, located at one of the five box positions (target box), and a visual memory period after which the array of boxes is presented again (reach object), randomly to the left or right of the screen center. During the following delay (movement planning period) the monkey has to maintain ocular fixation and withhold arm movement, thereby keeping body-, hand- and gaze-centered frames of reference aligned with the screen center. Disappearance of the hand fixation stimulus (Go signal) permits the monkey to reach to the memorized target box. The target box is defined by the position of the box that was cued on the reference object in object coordinates. Time spans of individual trial periods as depicted in the labels indicate the range of uniformly distributed random durations. **(b) Position invariance (Exp-I)**. Reference and reach object each are presented at fixed eccentricities randomly to the left and right of the screen center. Left and right offsets are uncorrelated, making the position of the reach object unpredictable and in 50% congruent and 50% incongruent to the reference object. The horizontal offset between left and right object center corresponds to the inter-box distance, so that left and right object location overlap in four boxes. This results in a total of six possible egocentric cue/reach target locations on the screen (Supplementary Fig 1). In Exp-I, reference and reach objects are always the same size and also otherwise visually identical. **(c) Size-invariance (Exp-II)**. In half of the trials the reference object had the same length as in Exp-I (example “long left” in upper row left) and in the other trials was half as long (example “short left” in upper row right). Again, reference and reach object each were presented randomly to the left or right of the screen center (Supplementary Fig 1). In Exp-II, horizontal offsets of reference and reach object differed in size and an additional vertical offset was introduced such that reference and reach object were incongruent in all trials (see Methods for details).

The animals memorized the location of a briefly flashed peripheral visual cue at one of five positions on an elongated visual object (**Fig. 1a**; reference object) to later reach towards this on-the-object position irrespective of object location on the screen (reach object). The egocentric cue positions varied independently of their object-centered positions since the reference object was presented with a left or right offset relative to the screen center, randomly in each trial (**Fig. 1b**). After a first delay period (visual memory), the object re-appeared as reach object, again randomly with a left or right offset. After a second delay (reach planning), the monkeys had to reach to the previously cued target position on the object. The location of the reference object was not predictive for the location of the reach object. In experiment I (Exp-I; **Fig. 1b**), reference and reach object were visually identical but their location could be shifted to test for position-invariant (=object-centered) encoding. In experiment II (Exp-II, **Fig. 1c**), additionally, the size could vary between reference and reach objects to test for position- and scale-invariant object-centered encoding (see Methods for details).

### Object-centered vs egocentric encoding hypothesis

In Exp-I, we asked if the fronto-parietal network encodes the location of the cue (visual memory) and the associated reach goal (reach planning) either in an object-centered or egocentric reference frame. **Fig. 2a** illustrates reference-object-left and reference-object-right selectivity profiles for two hypothetical neurons, representing object-centered and egocentric reference frames, respectively. In object-centered encoding, spatial selectivity of the neuron only depends on the position relative to the object. Thus, such neuron would show the same pattern of selectivity to different boxes on the object in object-left and -right conditions. Consequently, in body-centered screen coordinates (**Fig. 2a**, left) this corresponds to a shift of the selectivity profile that matches the object’s shift, while the shape of the profile stays the same.

**Figure 2:**
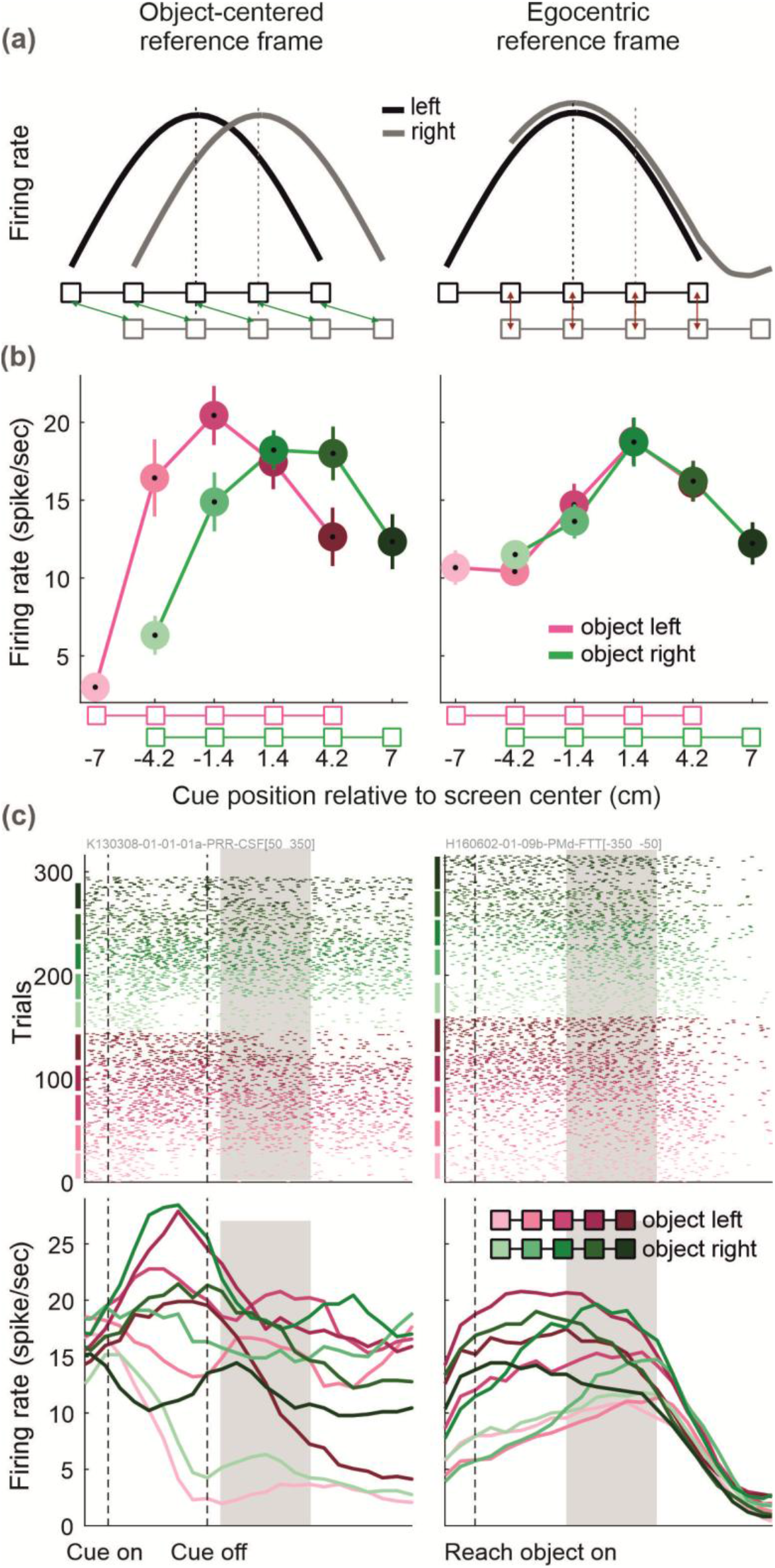
Object-centered and egocentric reference frames. **(a) Hypothetical object-centered and egocentric single unit responses.** The object-centered hypothesis (left column) predicts that a neuron keeps the same selectivity profile for different cue/target locations on the object, independently of the object location on the screen (=relative to the body). When analyzed as function of screen position, this would result in a shift of the profile together with the object. The egocentric hypothesis (right column) predicts that the selectivity profile is a function of the cue/target location relative to the body and not relative to the object. When comparing object-left and object-right profiles, the neuron shows the same response level for target positions overlapping in screen space, but different activity for non-overlapping boxes. Shifting of the object in this case would mean sampling a different part of the egocentric selectivity profile. **(b) Examples of single unit selectivity profiles from Exp-I.** An example single unit shows object-centered encoding (left column) of the cue during early visual memory period, 50 to 350 ms after cue offset (shaded time window in (c)). Another example shows egocentric encoding (right column) of the target during planning period, 350 to 50 ms before movement onset. The curves show mean firing rate across same-condition trials, error bars indicate SEM. Light/dark color shades correspond to the boxes on the object). **(c) Raster plot and PSTH of example units.** Left and right columns show raster plots (top row) and peri-stimulus time histograms (PSTHs, bottom row) from respective example units in (b). In the raster plot, every row is one trial and trials are sorted and grouped according to the on-the-object positions. The PSTHs below show spike frequencies (Gaussian kernel smoothing with 50 ms standard deviation) averaged across trials with identical object and on-the-object cue/target positions.

On the other hand, if an ideal neuron encodes the cue in an egocentric coordinate system, the selectivity will only depend on the egocentric location of the cue regardless of the object location. By shifting the object on the screen, one would sample different segments of an egocentric selectivity profile. Therefore, when comparing the object-left and object–right selectivity profiles, an egocentric neuron would show the same activity for positions with corresponding egocentric locations (**Fig. 2a**, right). This logic applies to cue encoding relative to reference object and reach goal encoding relative to the reach object.

### Single units in PRR and PMd encode location of the cue and reach goal in object-centered as well as in egocentric reference frames

Monkeys K and H performed the task with 72.15% ± 1.00% and 87.92% ± 1.20% average success rate across sessions. We recorded 100 and 107 single units in PRR and PMd, respectively, from monkey K, and 56 and 57 in PRR and PMd, respectively, from monkey H. From monkey K, 71 (71%) of neurons in PRR and 77 (72%) in PMd were included in the following analysis, from monkey H, 30 (54%) in PRR and 32 (56%) in PMd (see Methods). Since data from the two monkeys yielded corresponding results, throughout, we report the result from combining the two monkeys’ data unless mentioned otherwise.

During the visual memory period, selectivity profiles of a subset of cells in both areas were consistent with encoding of the cue location in an object-centered reference frame **(Fig. 2b-c**, left panels). The selectivity profile of this example neuron is independent of the screen-location of the object, hence object-centered. Single units with object-centered selectivity were found in both areas PRR (shown) and PMd of both monkeys.

Selectivity profiles of other units in the same areas and in both animals resembled egocentric encoding of the cue location (**Fig. 2b-c**, right panels). This example unit showed identical neural response strengths when overlapping boxes (i.e. egocentrically corresponding cue location) were cued with the object being located either left or right. Neural response strengths differed for non-overlapping boxes, i.e. when new locations relative to the body of the animal were sampled due to the new object location.

During the movement planning period, we also observed that selectivity profiles of some neurons resembled encoding of the reach goal in the object-centered reference frame and others in an egocentric reference frame (data not shown). As is not surprising for mixed encoding, during both the visual memory period and reach planning, selectivity patterns of most neurons only partly match the idealized patterns shown in **Fig. 2a**. Often neurons show more complex patterns with ambiguous or mixed object-centered and egocentric selectivity. Simple examples of ambiguity are linear selectivity profiles, in which case object-left and object-right tuning can result either from a horizontal shift or a vertical (firing rate) offset between both conditions (not shown).

To respect the continuous spectrum reflected in the observed mixed selectivity not only across areas or units, but also within individual units, we did not attempt to categorize the spatial selectivity of single units as binary object-centered or egocentric classes. Yet, the existence and gradual tendencies for either encoding scheme at the neural population level nevertheless can be quantified and might differ between brain areas or cognitive states and be informative about underlying computations.

### Predominant reference frame changes from object-centered during visual memory to egocentric during motor planning in PRR and PMd

During the late visual memory period, object-centered encoding of the cue location predominated across the neural population in both PMd and PRR. We quantified population-level predominance of either encoding scheme over the course of the trial in two ways: first, by computing a position invariance (PI) index, which is based on correlations of spatial selectivity profiles between conditions with different object positions (see Methods); second, with population decoding based on a cross-conditional classifier.

Positive values of the average (across neurons) PI during the memory period indicate predominant object-centered encoding (**Fig. 3a**). Once the location of the reach object is provided, the egocentric reach goal location becomes known to the monkeys for movement planning. After this reach object onset, the average PI shifted within 300-500ms, from predominant object-centered to predominant egocentric spatial selectivity in both brain areas (**Fig. 3a**). The single unit analysis allows to test if the shift of the PI distribution at the population level is due to selective activity of two different neuronal subpopulations in memory and planning periods (“recruitment”), or rather the consequence of a consistent shift in preferred encoding of the individual neurons (“re-coding”). When comparing visual memory and movement planning periods (**Fig. 3b**), the PI values cluster in the lower right quadrant and the angular density distribution is unimodal, as expected for the re-coding hypothesis. There is no indication of a clustering along the main axes and a corresponding bimodal angular density distribution, as would be expected for the recruitment hypothesis (Hartigan’s dip test; PRR p=0.84; PMd p=0.85). This re-coding pattern suggests that in PRR and PMd, the population of neurons contributing to either predominant reference frame in the two different periods mostly overlap. There is a weak but statistically significant negative correlation between PI of neurons in late memory period and movement planning period in PRR but not PMd (Pearson’s correlation coefficient; PRR r_p_ = −0.24, p = 0.015, PMd p = 0.3). This suggests that at least in PRR, on average, units with stronger object-centered preference in late memory tend to be more strongly egocentric in the later planning period.

**Figure 3.**
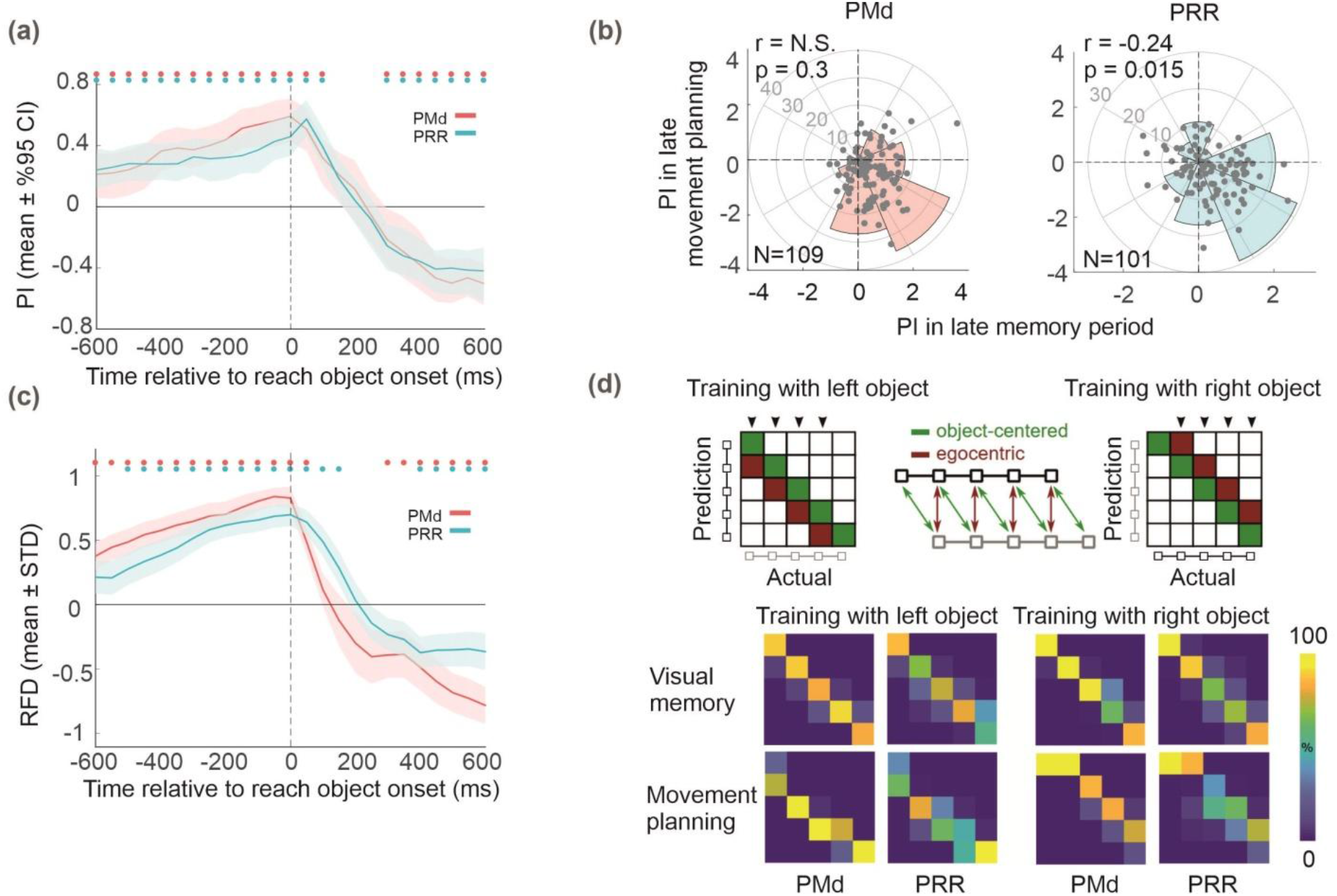
Population response with varying object position (Exp-I) **(a) Preferred reference frame across neuronal population.** Position Invariance (PI) measure across the population of 101 PRR (blue) and 109 PMd (red) single units (300 ms time bins, sliding by 50 ms). Positive and negative PI values indicate dominant object-centered and egocentric encoding, respectively. In the visual memory period (before reach object onset), object-centered encoding gained higher weight across the population, whereas reach goal locations (after reach object onset) were dominantly encoded in an egocentric reference frame. Dots at the top indicate significant deviation from zero (t-test, p values corrected for multiple comparison across time bins using False Discovery Rate correction (see Methods)) **(b) Reference frame in memory vs movement planning period.** PI values of individual units (dots in scatter plot) in late memory (300 ms before reach object onset) vs late reach planning period (300–600 ms after reach object onset) are shown. Angular densities across the population of neurons do not indicate deviations from a unimodal distribution (Hartigan’s dip test: PRR p=0.84; PMd p=0.85). r and p indicate Pearson’s correlation coefficient and its significance level. **(c) Relative difference of reference frame decoding (RFD).** Positive and negative RFD values indicate dominant classification of object-centered and egocentric positions, respectively similar to the time courses from the tuning analysis in (a). Dots at the top indicate significant deviation from zerp; p values corrected for multiple comparison across time bins using False Discovery Rate correction (see Methods). **(d) Hypothetical and actual confusion matrices.** In RFD, a decoder (5-way classifier) trained with the trials with the object on one side of the screen that is able to accurately classify the object-centered position of the target box when the object is on the other side of the screen. In the hypothetical confusion matrix (top row) these trials fall on the main diagonal (green). An egocentric encoding, instead, would introduce a systematic misclassification of the data, resulting in diagonals that are one off the main diagonal to the left or right (red), depending on which side is used for training the classifier. Arrows on top of the matrix mark the task conditions that are included in the calculation of the RFD values, as they have a distinct egocentric and object-centered representation (see Methods). Bottom row shows actual confusion matrices. During the late memory period object-centered positions are classified accurately. During late movement planning, a shift off the diagonal indicates classification of egocentric positions in both areas.

Second, we used neural population decoding to quantify the predominance of either reference frame. For this, we applied cross-conditional classification, and characterized generalization errors. Classifiers were trained to distinguish the five positions on the object using only trials where the object was on one side and tested its performance on trials where the object was on the other side. Predominant egocentric or object-centered neural reference frames, respectively, then predict characteristic classification patterns. High classification accuracy suggests an object-centered population encoding since each position on the object is decoded correctly irrespective of the object location relative to the body. Egocentric encoding instead would introduce systematic misclassification, namely a shift by one position off the main diagonal in the confusion matrix (**Fig. 3d**, top row).

From monkey K, 83-84 (83-84%) of neurons in PRR and 99-101 (93-94%) in PMd were included in the decoding analysis based on the number of available trials per condition (see Methods), from monkey H, 39-41 (70-73%) in PRR and 43-45 (75-79%) in PMd. There is a small variability in the number of neurons between conditions because the exact number of available trials depends on the condition that was used for training the decoder. When tested on the same data as used for training (iso-conditional), the classifiers showed high cross-validated classification accuracy of 95% for PMd and 80% for PRR on average over time bins and trial subsamples (data not shown). To test for the predominant reference frame, we computed the difference between the percentage of test trials classified in accordance with the object-centered hypothesis and the percentage classified in accordance with the egocentric hypothesis during the cross-conditional decoding (reference frame difference RFD; see Methods). The RFD (**Fig. 3c**) and confusion matrices (**Fig. 3d**, bottom row) show that during the visual memory period the cue position is predominantly classified in accordance with an object-centered reference frame (RFD > 0; PRR p < 0.01, PMd p < 0.01). Then there is a transition to classifying positions predominantly in an egocentric reference frame (RFD < 0; PRR p < 0.01, PMd p < 0.01) in the late movement planning phase.

### Single units scale the width of their selectivity profile with object size in PRR and PMd

Our results from Exp-I suggest a predominant object-centered encoding of the cue location during the memory period across the neuronal population. In Exp-II, we asked if the fronto-parietal network can encode on-the-object locations in an object-centered manner also for variable object size. Real-life objects might change their position not just within a fronto-parallel plane, but also in depth, thereby changing their visual size; or, they might exist in different-size variants, like hammers for different purposes. Hence, ideal object-centered encoding of on-the-object locations predicts that selectivity profiles of neurons should be characterized only by the relative positioning of the cue and the object, irrespective of object location (origin) and object size (scale), i.e., by position-invariant and size-invariant object-centered selectivity. In Exp-II, the monkeys saw a long or a short reference object, in randomly interleaved trials, relative to which they had to memorize the cue location (**Fig. 1c**). In order to preserve object-centered position information, one would expect a compression of the selectivity profile (in the egocentric screen space) for object-short compared to object-long trials (**Fig. 4a**, left). In contrast, egocentric encoding would result in sampling of a reduced range of the selectivity profile in object-short trials (**Fig. 4a**, right).

**Figure 4.**
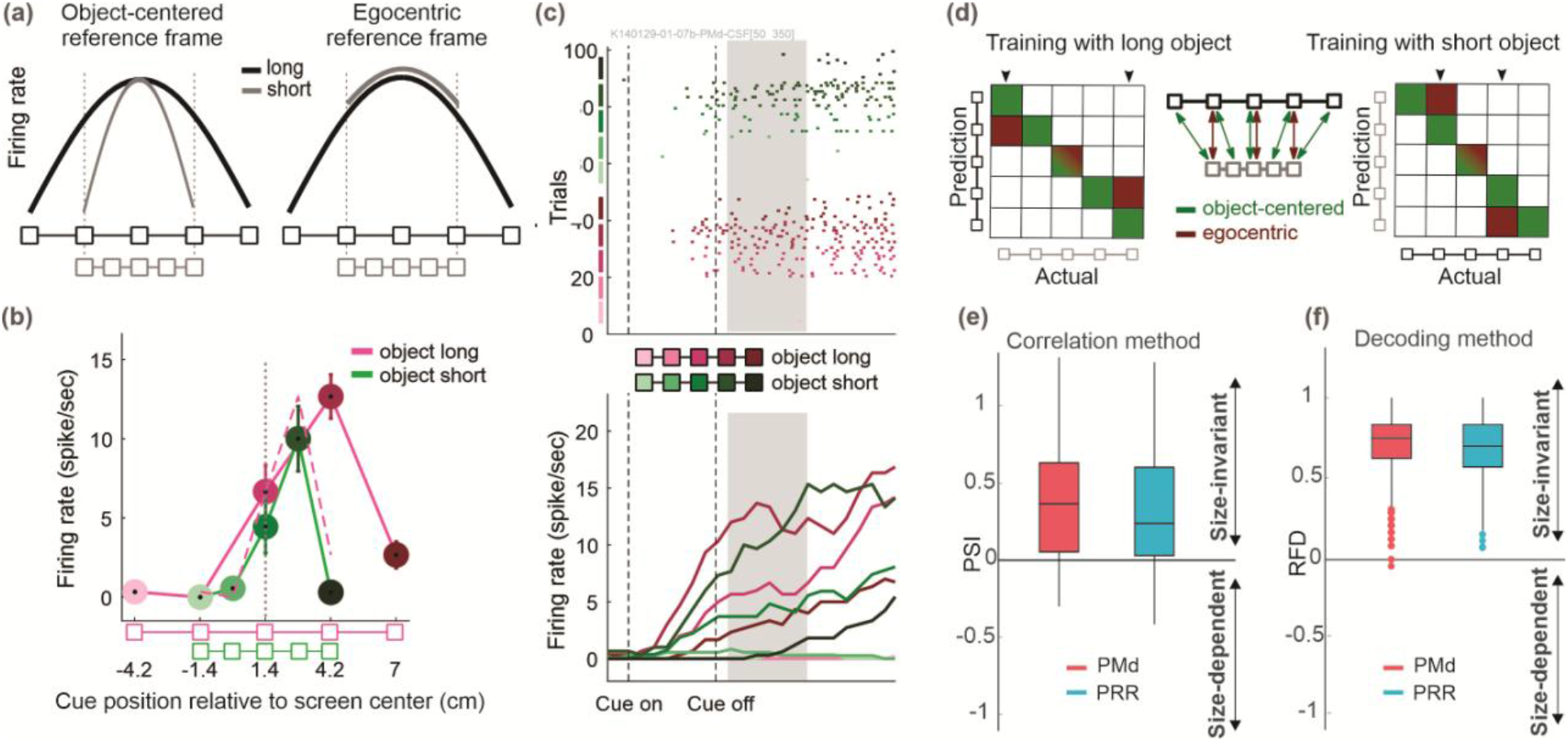
Size-invariant object-centered encoding (Exp-II) **(a) Hypothetical size-dependent and -invariant single unit responses.** Selectivity profiles of ideal object-centered size-invariant (left) and egocentric size-dependent (right) neuronal encoding. The size-invariance hypothesis predicts that the selectivity profile scales with the object size, such that the profile to different locations on the object is the same irrespective of object size. The egocentric hypothesis predicts the same response level for positions with identical egocentric locations, thereby sampling less horizontal extend when objects are small. **(b-c) Example unit from Exp-II.** Activity of an example unit when short and long reference objects were presented at the right side of the screen center. Object-centered size-invariant encoding is identified by a horizontally scaled selectivity profile. The dashed curve represents the ideal object-centered scaling of the selectivity profile for short-object trials, predicted from the selectivity profile observed in long-object trials, assuming that the width of the activity profile was scaled with the size of the object and relative to center of the object. The actually measured profile in this example precisely matched the prediction except for a small reduction in response gain by about 10%. If normalizing to the maximal response, result and prediction match exactly (not shown). All other settings are as in **Fig. 2b-c**. **(d) Hypothetical confusion matrices.** In RFD, a decoder (5-way classifier) trained with the trials with one object size that is able to accurately classify the object-centered position of the target box when the object is of the other size. In the hypothetical confusion matrix these trials fall on the main diagonal (green). An egocentric encoding, instead, would introduce a systematic misclassification of the data, resulting in diagonals that are compressed compared to the main diagonal, vertically or horizontally (red), depending on which object size is used for training the classifier. Arrows on top of the matrix mark the task conditions that are included in the calculation of the RFD values, as they have a distinct egocentric and object-centered representation (see Methods). **(e)** Mostly positive Position and Size Invariance (PSI, see Method) values across the population of 66 PMd (red) and 70 PRR (blue) single units indicate predominant size-invariant neural selectivity in the last 300 ms of visual memory period (signed-rank test, PRR p < 7×10^-9^, PMd p < 10^-9^). **(f)** Equivalently, positive RFD values indicate dominant classification of object-centered rather than egocentric positions, respectively. The box plots show median and the 75^th^ (top) and 25^th^ (bottom) percentiles, as well as the data range (whiskers)without putative outliers (dots; more distant from 25/75 percentiles than 1.5 times the respective interquartile range).

For Exp-II, we recorded 67 and 53 single units in PRR and PMd, respectively, from monkey K; 36 and 45 single units in PRR and PMd from monkey H. We identified 51 (76%) and 46 (87%) active units in PRR and PMd from monkey K, 19 (53%) and 20 (44%) from monkey H, which were included in the following analysis. Monkeys K and H performed the task with 63.94% ± 0.88% and 81.70% ± 2.50% average success rate.

Neurons in PRR and in PMd of both monkeys revealed examples of size-invariant object-centered encoding during visual memory. **Fig. 4b** and **4c** show an example unit where the object-short response profile fits the prediction of the object-centered hypothesis in the sense that it represents a duplicate of the long-object selectivity profile compressed by the factor that short and long object differ in length. The example neuron showed the same selectivity to long and short reference objects, respectively, when they were presented on the left of the screen center (not shown). In other words, only the relative on-object position determined the response, not the location of the object on the screen nor the size of the object. As in Exp-I, also in Exp-II we observed units with more complex selectivity profiles which either do not or only partly fit the object-centered or the egocentric hypothesis (**Supplementary Fig. 2**). We tested for size invariance only during the visual memory period, not during movement planning, since Exp-I had shown dominant object-centered encoding during the memory period only. Consequently, in Exp-II, short objects were only used as reference objects, not as reach objects.

In order to assess the contribution of position- and size-invariant encoding at the neural population level, we again computed the correlation measure and the decoding measure. For the correlation measure, we computed PSI for each neurons selectivity profiles (see Methods). The distribution of PSI across units (**Fig. 4e**) shows that different units scale their selectivity profile to different extent, evident from the range of PSI values across the population. However, on average, size-invariance predominated (signed-rank test, PRR p = 7×10^-9^, PMd p = 10^-9^), consistently across areas.

With the cross-conditional classification approach, we tested for generalization of the decoder between the long and short reference object conditions (see Methods). The cross-validated accuracy of the classifiers on the training data was high during the visual memory period, 88% for PMd and 80% for PRR on average over time and repetitions. The RFD in **Fig. 4f** shows that in both areas the cue position predominantly can be decoded invariant to the size of the object during the late visual memory period (RFD > 0; PRR p < 0.01, PMd p < 0.01).

## Discussion

Reaching towards and grasping objects is a major way in which primates interact with their environment. Object-relative position information is important in guiding these movements. Such allocentric alongside egocentric frames of reference provide a stable representation of space which is robust to environmental dynamics and noise (Burgess, 2006). Having such stability in spatial representations is crucial for goal-directed reach planning when, for example, the reach goal is located on a moving object or when the object is temporarily occluded. Here we showed, first, that object-centered spatial encoding is part of the mixed ego- and allocentric neural selectivity found in the frontoparietal reach planning network when reach is directed towards different on-the-object sites. Second, in both, PRR and PMd the predominant frame of reference shifts according to cognitive demands in different epochs of the behavioral task. During visual memory, when the cue-on-object location was the most parsimonious way of maintaining the relevant spatial information, object-centered encoding predominated. During movement planning, when the actual reach goal could be determined from the integration of the memorized cue-on-object location and the now visible and stationary reach object location, egocentric encoding predominated. Third, by providing evidence for size-invariant spatial selectivity, we demonstrate that allocentric encoding not only means that the origin of the reference frame is centered on the object, but also that the spatial scale scales with object-size. Such size-invariant representation of object-related information so far was associated with object-recognition tasks and mainly ventral stream processing (DiCarlo and Cox, 2007; Grill-Spector and Weiner, 2014; Tacchetti et al., 2018), but not with dorsal stream movement preparation.

For successful physical interactions, a geometrical representation of the object, including features such as object size and on-the-object sites, is necessary for action planning. Additional to pre-shaping the hand to fit the geometry of the object before grasping (Aglioti et al., 1995), a proper object-relative placement of the hand is needed, e.g., to pick up a hammer at its handle instead of its head. Correct object-relative hand positioning is relevant for successful completion of the object-associated action, and hence might differ from allocentric localization of reach goals relative to a landmark with which one is not going to have direct physical interaction. When localizing reach goals relative to a landmark, human imaging data suggest that spatial processing along the dorsal stream is predominantly egocentric (Chen et al., 2014). In contrast, when human subjects had to spatially judge reach goals based on geometrical features like relative distance and direction of a remote pair of dots, the frontoparietal network was more active for such allocentric movements compared to egocentric movements directed towards a visual target (Thaler and Goodale, 2011c).

We see that either reference frame can dominate in the frontoparietal reach network. The allocentric frame dominated while the animals had to memorize an on-the-object position to generalize this geometric information to the time when the final object location on the screen would become available. The egocentric frame dominated as soon as animals could plan an according reach movement (Chen et al., 2014). While mixed egocentric reference frames have been described before (Chang and Snyder, 2010; McGuire and Sabes, 2011), our findings further suggest that predominant reference frames are not fixed properties in frontoparietal association cortices. Instead, they reflect task-specific computations to achieve spatial transformation, thereby not just giving variable weight to different egocentric sensory modalities (Bernier and Grafton, 2010; Bremner and Andersen, 2014), but also including allocentric computations if demanded by the task. The diversity of preference, which ranges from egocentric to object-centered encoding across neurons in PRR and PMd provides high flexibility for the fronto-parietal network for conveying spatial information to recipient areas in either of the two frames of reference.

Overall, human imaging studies suggest partially overlapping networks for allocentric reach movement planning (target memory) and egocentric guidance of movements. Yet, while (Chen et al., 2018, 2014) reported strongest allocentric preference in temporal cortices, they attribute mixed ego- and allocentric encoding for computing allo-to-ego conversions to human dorsal premotor cortex (PMd), pre-supplementary motor area (pre-SMA) and precuneus, but not to medial intraparietal sulcus (mIPS), the superior parietal lobe (SPL) or the superior parieto-occipital (SPOC), which were predominantly egocentric (Chen et al., 2014; Chen and Crawford, 2020). Our data provides details at the single neuron level about mixed encoding and the transition from predominant allocentric encoding to egocentric encoding as soon as the reach goal is determined (Byrne et al., 2010), corroborating the predictions for dorsal premotor cortex from the human neuroimaging studies (Chen et al., 2018). Yet, different to what human imaging suggests, we see a marked similarity between PRR and PMd regarding allocentric encoding.

The relevance of allocentric, especially object-centered encoding in skeletomotor compared to oculomotor tasks was unclear so far, since detailed monkey neurophysiology data existed only for saccade tasks. The importance of object geometry for manual interaction with objects may explain why we found allocentric encoding while neurons in the lateral intraparietal area (LIP) did not show predominant object-centered encoding during an object-relative saccade task, which was otherwise similar to our task (Sabes et al., 2002). On the other hand, saccade studies in monkey parietal area 7a suggest that individual neurons can encode target location not only in gaze-centered (egocentric) reference frame (Andersen et al., 1985), but also show left-right selectivity for spatial information relative to task-relevant objects (Chafee et al., 2007; Crowe et al., 2008). Similarly, subsets of neurons in the supplementary eye fields (SEF) of the frontal cortex have been shown to be selective for the left or right end of an object or a pair of dots towards which a saccade is directed (Moorman and Olson, 2007; Olson et al., 2000; Olson and Gettner, 1999, 1995; Olson and Tremblay, 2000).

Yet, for binary left-right selectivity, it can be difficult to distinguish categorical rule-like encoding from an object-relative position code proper. Given that neurons, especially in the frontal areas, are known to represent categorical abstract rules (Wallis et al., 2001; Wallis and Miller, 2003), it was speculated if object-relative left-right neural selectivity could result from top-down rule signals in previous object-centered saccade experiments (Filimon, 2015). In our task, we sampled space-continuous selectivity profiles with five on-the-object positions. Rule encoding would predict identical selectivity profiles for object left/right/long/short conditions, which was not the case for most of the units we recorded. Instead, we observed a spectrum of ego- and object-centered encoding, including mixed reference frames and partial scaling, as typical for the computation of coordinate transformations in neural networks (Avillac et al., 2005; Brozovic et al., 2007; Deneve et al., 2001; Deneve and Pouget, 2003), which cannot be explained merely by rule encoding. We therefore interpret our observed neural selectivity profiles as indication of spatial encoding supporting the transformation between ego- and allocentric representations of space in the context of skeletomotor planning, rather than categorical rule encoding.

Remarkably, object-centered encoding was accompanied with size-invariant spatial encoding of different on-the-object sites, at the single neuron and population levels. To our knowledge, this is the first electrophysiology evidence showing that the fronto-parietal network utilizes size-invariant positional code for movement planning.

Size-invariance is a coding property discussed in the context of object recognition and associated with ventral stream processing (DiCarlo and Cox, 2007; Grill-Spector and Weiner, 2014; Tacchetti et al., 2018). Yet, the relative position of object features is not just relevant for recognition, but also for interaction with the object, e.g., to pick up different sized hammers always at the end of their handle. Since we presented reference array and cue not simultaneously (in one monkey), we consider it unlikely that the animal memorized the position information by means of a visual pattern and that the size-invariant selectivity observed here “echoes” ventral stream pattern encoding.

Previous studies showed generalized quantity encoding in neurons in the depth of the intraparietal sulcus of monkeys (Tudusciuc and Nieder, 2007) and in parietal BOLD signals in humans (Harvey et al., 2015), i.e., selectivity for quantity irrespective if quantity was presented via the size of an object or the number of visual items. Due to the discrete nature of the on-the-object cue positions, it could be that the animals memorized them via an object-relative (left-most, middle,…) or an abstracted positional code (1^st^, 2^nd^, 3^rd^,… position on the object) which is numerical but still object-related. Translating back such abstracted numerical information into spatial motor goal information would be a highly useful capacity, e.g., for foraging in discretized environments (“5^th^ tree from the left”).

PRR and PMd are along the course of the dorsal visual processing pathway and we showed that they express allocentric and size-invariant encoding with a short latency of a few hundred milliseconds after presentation of an object-relative spatial cue. Allocentric and size invariant abstract representations are two important properties which according to the classical two visual pathways model (Goodale et al., 2004; Goodale and Westwood, 2004; Milner and Goodale, 2008) are associated with the areas along the ventral stream. The model suggests that in memory-guided actions, the dorsal stream could access the ventral stream neural codes through interaction with different areas in the ventral stream. However, it does not predict the latencies of the interaction and how quickly the codes could be accessible in the dorsal stream. If the model accounts for our results, as we had predicted in our earlier behavioral study (Taghizadeh and Gail, 2014), our findings strongly suggests that the interaction across the two pathways are almost immediate and does not take the long latencies which had been previously suggested by behavioral experiments (see (Westwood and Goodale, 2011) for review). Functional and direct and indirect anatomical connections between frontal and parietal cortex with areas of the ventral stream exist ((Borra et al., 2010), see reviews from (Cloutman, 2013; Perry and Fallah, 2014)). Our data suggest that in order to plan movements which take higher levels of computations and cognitive load for interacting with objects according to geometrical considerations, the two processing streams need to be functionally tightly connected and work together as a network to support the dorsal stream for flexible action planning to interact with non-stationary dynamic environments.

## Methods

### Object-directed memory-guided reaching task

The monkeys were seated in a primate chair in a dimly lit room in front of a fronto-parallel touchscreen. With the help of head-fixation and trained gaze fixation (see below), the monkeys’ mid-sagittal plane and all egocentric references were aligned to the screen center. Visual stimuli were presented on an LCD screen (19” ViewSonic VX922; onset latencies corrected; background intensity of 0.16 cd/m2) mounted behind the touchscreen (IntelliTouch, ELO Systems, CA, USA). The distance between the monkeys’ eyes and the screen was 39-45 cm. Throughout the text, conversions from centimeter to degree are based on a 40 cm distance.

The temporal structure of the task was identical in Exp-I and II (**Fig. 1a**). The monkey initiated a trial by acquiring central gaze fixation (224 Hz CCD camera, ET-49B, Thomas Recording) and hand fixation on the touchscreen. The gaze fixation stimulus was a filled red square of 0.5 cm (0.72°) side length and 7 cd/m^2^ intensity, and the hand fixation stimulus was a filled white square of 0.5 cm (0.72°) side length and 13 cd/m^2^ intensity. Gaze and hand fixation was enforced within 2-3 cm (2.86-2.89°) around each of the two immediately adjacent fixation points. In case of unsuccessful fixation, the trial was aborted and repeated at a random later time during the experiment.

Valid gaze and hand fixation for a random fixation period of 600-1100 ms was followed by a 400ms presentation of an array of five boxes, horizontally arranged and connected with a line (reference object; details see below). The boxes indicated possible positions of the upcoming spatial cue. The cue consisted of a small dot of 0.27 cm (0.39°) diameter presented at the position of one of the five reference object boxes. To balance task conditions, cue positions were selected pseudo-randomly with increasing probability of so-far underrepresented conditions, and such that the difference in number of correct repetitions between different positions did not exceed three trials.

After an inter-stimulus interval (ISI) of 80 ms following the offset of the reference object, a spatial cue was presented for 340 ms. For monkey H, the ISI was removed and the reference object remained visible during the 420 (=80+340)ms of cue presentation time. The cue presentation was followed by a first variable memory period of 700-1200 ms for monkey K and 900-1200 ms for monkey H, during which only the fixation stimuli were shown (visual memory). After this delay, a reach object of the same type as the reference object was presented. The monkey was instructed to later touch the box on the reach object which corresponded to the box that was cued on the reference object, e.g., for a cue seen at the left-most box of the reference object, the monkey should reach towards the left-most box of the reach object (target), irrespective of the absolute position of the reach object on the screen. The onset of the reach object was followed by a second delay period of 615-646 ms during which the reach object was visible but the monkey was required to maintain gaze fixation and to withhold the movement (movement planning). Continued visibility of the reach object allowed the animals mentally maintaining the reach goal location either in an object-centered or egocentric reference frame. After the second delay period, the hand fixation stimulus disappeared. This served as the GO cue to reach to and touch the cued on-the-object target location within 1000 ms while holding central ocular fixation. Reach endpoints had to be within an elliptical area (horizontal semi-minor axis 1.2 cm (1.72°); vertical semi-major axis 4 cm (5.71°)) around the target box. After holding the target for 220 ms (monkey H; 300ms monkey K) the trial counted as successful and the monkey received a visual (a small, light gray dot of 0.27 cm (0.39°) diameter on the target box of the reach object), an acoustic (a high-pitched tone) feedback, and a drop of juice as reward. The monkey’s mid-sagittal plane, gaze and hand fixation points aligned to the center of the screen. The reference object randomly presented at one of two egocentric locations, left or right of the screen center, with equal eccentricity (**Fig. 1b**). The reach object also presented either left or right, but the possible locations differed across experiments I and II (see below). Here and throughout the text, “location” of the object refers to the position of the center of mass of the object on the screen, i.e., the center point of the central box on the object. Reference object, reach object and cue had a low intensity gray tone (2 cd/m^2^).

#### Exp-1 Position invariance

In Exp-I, reference and reach objects where always identical in terms of visual appearance. The individual boxes of the reference and reach objects were 0.35 cm (0.50°) squares with 2.8 cm (4.00°) center-to-center distance. In terms of location, reference and reach object either matched (position-congruent trials) or differed (position-incongruent). Left and right screen locations were (x,y) = (±1.4, 2.7) cm = (±2.0, 3.8) ° relative to screen center. This means, objects were vertically elevated above the eye and hand fixation position to prevent visual interference with fixation stimuli and obstruction by the animal’s arm. Horizontally, in position-incongruent trials, the locations of the reference and reach object boxes were set off by one box distance, such that four out of five box positions overlapped between reference (potential cue locations) and reach object (potential reach targets). For example, boxes 2 to 5 (counting from left to right) of a left-side object had identical egocentric locations to boxes 1 to 4 of a right–side object (**Fig. 1b**). Therefore, while the cue (target) could take five different positions relative to the reference (decision) object, i.e., five different object-centered positions, they covered in total six different potential egocentric locations on the screen. The 20 different combinations of cue, reference and reach object positions (5×2×2) were presented pseudo-randomly (algorithm as above). By the nature of the behavioral task, the cue and the reach goal always had the same object-centered position. In position-congruent trials, the cue and the reach goal additionally had the same egocentric locations. In position-incongruent trials, the cue and the reach goal differed in egocentric position and the reach goal needed to be determined based on the object-based location of the target box. Since the congruency of the trials was unpredictable, the monkey was encouraged to follow object-centered encoding of the cue in all trials for successful task performance. The monkeys could only determine the egocentric reach goal location upon occurrence of the reach object.

#### Exp-2-Size invariance

In Exp-II, reference and reach objects always differed in terms of location (position-incongruent trials only) and could additionally differ in size. In long-object trials (size-congruent, **Fig. 1c,** top left), the reference object was identical to Exp-I. In short-object trials (size-incongruent, **Fig. 1c,** top right), the boxes of the reference object were only 1.4 cm (2.00°) apart, i.e. half the spacing of the long object. The reach object in both long and short trials was identical to the long reference object. Both long and short reference objects were presented at the same left and right locations as in Exp-I. Unlike Exp-I, the reach object in Exp-II was always position-incongruent to the reference object. While the reference and reach object could be congruent or incongruent in terms of horizontal location (as in Exp-I), the reach object always had a vertical position offset and was located at (x,y) = (±2.3, 4.4) cm = (±3.4, 6.3)° (**Fig. 1c,** bottom). This offset was introduced to maximally encourage object-based encoding of the cue location during the visual memory period, since the purpose of Exp-II was to specifically test for scale invariance during object centered encoding. Again, by nature of the task, the cue and the reach goal always had the same object-centered position. Long- and short-object trials were presented in alternating blocks. In long-object blocks, as in Exp-I, all 20 different combinations of cue, reference and reach object positions (5×2×2) were pseudo-randomly presented. In short-object blocks, reference object left and right conditions were blocked for ease of performance. In each sub-block, the 5×2 combinations of cue and reach object locations where randomly interleaved such that horizontal (in-)congruency was always unpredictable and hence the reach goal unknown prior to onset of the reach object.

#### Behavioral success rate

was calculated as percentage of correctly performed reaches relative to initiated trials in which the monkey continued until after he started the reach movement.

#### Single unit selection

In both experiments, only “active” units were included in the reference frame analysis and were defined as units which on average (across all trials, regardless of task condition) fired at least 5 spikes within a time interval extending from 100 ms before until 1500 ms after cue onset in correctly performed trials. Average firing rate across all selected units in Exp-I for PRR and PMd was 15.63 and 15.02, respectively, in monkey K; 12.03 and 10.66 in monkey H. In Exp-II for PRR and PMd it was 19.1 and 15.7 in monkey K, and 10.6 and 11.6 in monkey H.

#### Correlation method for testing object-centered and egocentric reference frames hypotheses based on single units’ firing rates

At the single unit level, we distinguished object-centered and egocentric reference frames by comparing each neuron’s spatial selectivity profile for the five cue stimulus positions in trials when an object was presented on the left side with the trials when the corresponding object was presented on the right side. During visual memory, we compared the reference object positions, irrespective of reach object position. Vice versa, during movement planning we compared the reach object positions, irrespective of reference object position. The object-centered hypothesis predicts object-left and object-right selectivity profiles to be identical in shape, but shifted relative to each other in egocentric screen space (**Fig. 2a-c, left**). The egocentric hypothesis instead predicts the selectivity profiles to be identical only in the range of object compartments that overlap in egocentric screen space (**Fig. 2a-c, right**). For each cell, we measured the similarity of the two selectivity profiles to quantify how well either hypothesis can explain the response of the neuron. We used the Pearson’s linear correlation coefficient, similar to (Mullette-Gillman et al., 2005), between each pair of the selectivity profiles, i.e., reference object left versus right; reach object left versus right. For variance-stabilization, the bounded correlation coefficients were subjected to Fisher’s z transformation (inverse hyperbolic tangent function) before using them for any further analysis and statistical tests. For brevity, we refer to the Fisher z-transformed correlation coefficients as “correlation coefficients”.

Since the correlation coefficient is insensitive to linear scaling, the similarity between the shapes of the selectivity profiles is quantified independent of potential gain modulation effects on the neural firing rates. This means, if object location modulates responses to all cue positions by an equal factor, without affecting relative response strengths for different on-the-object cue positions, the correlation method considers this outcome to be in accordance with the object-centered hypothesis. Our complementary decoding method (see below) is sensitive to gain effects on neural activity.

For each cell and pairwise selectivity profile, the correlation coefficient was calculated twice, once in each reference frame. The *object-centered correlation (objCorr)* is the correlation coefficient between the samples of the object left and right selectivity profiles with corresponding object-centered cue locations. (**Fig. 2a**, left, green arrows show points with equivalent object-centered location). The *egocentric correlation (egoCorr)* is the correlation coefficient between samples of the object left and right tuning curves with corresponding egocentric locations. (**Fig. 2a**, right, dark red arrows show points with equivalent egocentric location). Each of the correlation values provides a measure that determines the validity of the corresponding hypothesis for each cell’s activity profile. The difference *objCorr-egoCorr* between them indicates the neuron’s preference of reference frame. We will refer to this difference as Position Invariance (PI). Positive values indicate selectivity profile of the neuron was better explained by the object-centered hypothesis (= invariance with respect to egocentric location), while negative values point towards the egocentric hypothesis (= invariance with respect to object-centered location).

In order to reduce effect of outliers in trial-by-trial firing rates, we estimated the mean firing rate for every unit with a bootstrap method (200x resampling; bootstrp() function in MATLAB) and the PI was calculated for every bootstrap sample. In addition, for every bootstrap sample, the null distribution of PI was estimated by randomly shuffling trials (trials used to generate this bootstrap sample) 100 times, across all task conditions, and the average null-PI was subtracted from the corresponding PI.

#### Visual memory vs movement planning reference frame

To study changes in the preferred reference frame across time, for each cell PI was calculated in 300 ms windows, sliding by 50 ms, extending from 600 ms before until 600 ms after reach object onset (data was aligned to reach object onset). In all time bins earlier than reach object onset (up to bin [-150 150] ms), selectivity profiles were computed for cue relative to reference object (memory period); after reach object onset, selectivity profiles were computed for target relative to reach object (movement planning period). The time bin centered on reach object onset ([-150 150]ms) was included in both memory and movement planning periods.

When comparing PI in visual memory versus movement planning, we tested across the population of neurons, if a strong preference for one reference frame during visual memory would be associated with a strong preference for the same, the other, or no preferred reference frame during movement planning. For this, we plotted the PI of each neuron in memory vs planning period on orthogonal axes (**Fig. 3b**), and measured the angular deviation θ from the x+ axis, between −180 and 180 degrees, corresponding to the arctan of the ratio PI_planning/PI_memory. Neurons showing a preferred reference frame selectively in only one delay period fall onto one of the main axes, neurons with strongly (anti-) correlated preference fall on the diagonals. Predominant subpopulations of either type would result in an inhomogeneous distribution of the angles. Distributions were estimated by binning the data into 45° bins, and subjected to Hartigan’s dip test for multimodality. We compared responses 300 ms preceding (late memory) and 300-600 ms after (late planning period) reach object onset, when the neural activity is comparably stable after the transient change in stimulus.

#### Scaling of the spatial selectivity to the object size in Exp-II

Exp-II aimed at further characterizing object-centered encoding. Therefore, we analyzed firing rates only during the late memory period when neurons have reached their sustained activity and predominant position-invariant (object-centered) preference was expected based on the results of Exp-I. The object-centered hypothesis predicts that selectivity profiles should scale with the object size in the egocentric screen space (**Fig. 4a**, left), independent of its location, meaning similar object-centered selectivity profiles in all four conditions of object size and location. The egocentric hypothesis predicts that selectivity profiles for small object size should resample part of the selectivity profile for large object size in the egocentric screen space (**Fig. 4a**, right). To test this, we quantified two correlation coefficients for every neuron: *alloCorr* was the correlation coefficient between a neuron’s activity in ten conditions (5 on-the-object cue positions times 2 reference object locations) in long-object trials and their corresponding conditions (with identical object-centered cue location) in short-object trials (2 sets of 10 green arrows in **Supplementary Fig. 1a**). *egoCorr* was the correlation coefficient between neuron’s activity in six conditions (3 on-object cue positions times 2 reference object locations) in long-object trials and their corresponding conditions (with identical egocentric locations) in short-object trials (2 sets of 6 red arrows in **Supplementary Fig. 1b**). To account for both size and position invariance at the same time, alloCorr and egoCorr were calculated across all possible pairing of the firing rates between the short and long trials, regardless of object location. We calculated *alloCorr-egoCorr* to quantify how far a neuron complies with the prediction of object-centered Position and Size Invariant (PSI) encoding, and refer to this measure as PSI. In the same way as in Exp-I, PSI for every cell was bootstrapped and a shuffle predictor subtracted.

#### Decoding population activity

As an additional population-level analysis, we used a cross-conditional decoding approach: A 5-way classifier for decoding cue or target position on the object was trained with data from one condition (e.g. reference object located left on screen) and its performance was tested on data from another condition (reference object right). For Exp-I, there are two conditions, object-left and object-right, in the memory and planning period, respectively. A decoder that is only trained on trials of the object-left condition but accurately predicts the position in object-right condition trials (left-right generalization) suggests a predominant position-invariant (object-centered) encoding in the data. This will show as a confusion matrix where only the main diagonal is populated (**Fig. 3d**). On the other hand, a systematic offset in the predicted labels, e.g., object-right trials at positions 1, 2, 3, 4 (counting the boxes from left to right) are decoded as positions 2, 3, 4, 5 by the object-left classifier, is indicative of an egocentric encoding, because these positions correspond to the same egocentric locations. Such pattern would show as a confusion matrix populated along the secondary diagonal. In Exp-II, there are four conditions, *long-object-left*, *long-object-right*, *short-object-left*, *short-object-right*, and each pair has to be compared independently. To quantify the size-invariance of the neural population, we compared each *long*-with each *short-object* condition, resulting in four comparisons.

For the decoding analysis, we included only units for which we recorded at least 15 trials per cue/target position in the training condition. This enabled us to select a random subset of 15 trials per position of the training condition from each unit. The average firing rates of the selected units in a 300ms window during the selected trials were combined to form the 75 feature vectors for training the decoder. Feature vectors for testing were built similarly by combining a random subset of as many trials as possible (same number for each position) of the testing condition from the same units. This randomization was repeated 1000 times, each time choosing random subsets of trials from each unit. This way, we can include the whole dataset in the decoding analysis, despite each unit being recorded for a different number of trials. For each repetition, an independent classifier was trained. We used an error-correcting output codes (ECOC) model with 10 binary linear support vector machines as the classifier (MATLAB function: fitcecoc). To get a time-continuous estimate, we repeated the analysis, each time shifting the window by 50ms. During the visual memory period, the condition of each trial was defined by the reference object, during the movement planning period it was defined by the reach object. For the time windows that overlapped with both periods (around reach object onset), the condition was defined by the period with the most overlap. For the window centered on the reach object onset, both definitions where used.

In order to quantify the dominance of a reference frame, we calculate the difference *p_obj_* – *p_ego_* in percentage of trials decoded in accordance with the object-centered hypothesis and percentage of trials decoded in accordance with the egocentric hypothesis. This measure will be referred to as Reference Frame Difference (RFD). A value of 1 shows complete object-centered encoding, while a value of −1 shows complete egocentric encoding. For this calculation, we only consider the test trials to positions that have an egocentric equivalent in the training condition. For example, we cannot include test trials to the right-most position when the object is on the right, because this location does not exist in the object-left condition (**Fig. 1a** and **Fig. 3d**). The number of eligible positions depends on the conditions that are being compared: In Exp-I, four positions overlap (**Fig. 3d**) while for comparisons of short and long objects in Exp-II, two positions can be considered (three positions overlap, but one of these is the same in both reference frames and hence does not help to distinguish them, **Supplementary Fig. 1**).

To test for a significant difference from zero, we calculate the RFD for each of the 1000 randomizations and compute the percentage of values lying below and above 0. If one of these percentages is less than half the alpha criterion, it is considered significant.

In all analysis where results of statistical tests were compared across multiple time bins, the significance level was adjusted to account for multiple comparisons by correcting for the false-discovery rate, allowing a proportion of false positives of less than 5%.

#### Animal implantation and neural recordings

The procedures for animal preparation and neural recordings were described previously (Westendorff et al., 2010) and are here repeated for completeness. Numerical values have been adjusted to the current experiment. Two monkeys implanted with a titanium head holder and two magnetic resonance imaging (MRI)-compatible recording chambers, custom-fit to the monkeys’ heads (3di, Jena Germany; and Thomas Recording, Giessen, Germany). Chamber positioning above PRR (Horsley Clarke coordinates: 10 mm contralateral and 13 mm posterior for monkey K; 12.5 mm contralateral and 13.5 mm posterior for monkey H) and PMd (18.5 mm contralateral and 20 mm anterior for monkey K; 19 mm contralateral and 22 mm anterior for monkey H) was guided by pre-surgical structural MRI and confirmed by postsurgical MRI. Sustained direction-selective neural responses during center-out reach planning (memory period) served as physiological signature to confirm the region of interest in both areas. Both chambers implanted contralaterally to the handedness of the monkey (left hemisphere). All surgical and imaging procedures were conducted under general anesthesia. We used two five-channel microdrives (“mini-matrix”; Thomas Recording) for extracellular recordings, mostly simultaneously in both chambers. The raw signals of the electrodes were pre-amplified (20x; Thomas Recording), band pass filtered, and amplified (154 Hz to 8.8 kHz; 400–800x; Plexon) before online spike sorting was conducted (Sort Client; Plexon). Spike times and spike waveforms were recorded and later subjected to additional offline sorting (Offline Sorter; Plexon).

Both animals were housed in social groups with one or two male conspecifics in facilities of the German Primate Center. The facilities provide cage sizes exceeding the requirements by German and European regulations, and access to an enriched environment including wooden structures and various toys. All procedures have been approved by the responsible regional government office [Niedersächsisches Landesamt für Verbraucherschutz und Lebensmittelsicherheit (LAVES)] under permit numbers 3392 42502-04-13/1100 and comply with German Law and the European Directive 2010/63/EU regulating use of animals in research.

## Acknowledgements

We thank Sina Plümer for help with data collection and technical support, Klaus Heisig for help with setup maintenance, Leonore Burchardt for help with animal training and Dirk Prüße for technical support.

This work was supported by and benefited from the State of Lower Saxony (grant VWZN2563), the European Commission in the context of the Plan4Act consortium (http://plan4act-project.eu; EC-H2020-FETPROACT-16732266), and the German Research Foundation in the context of the Collaborative Research Center Cognition of Interaction (DFG SFB-1528).

---

**Supplementary Figure 1.**
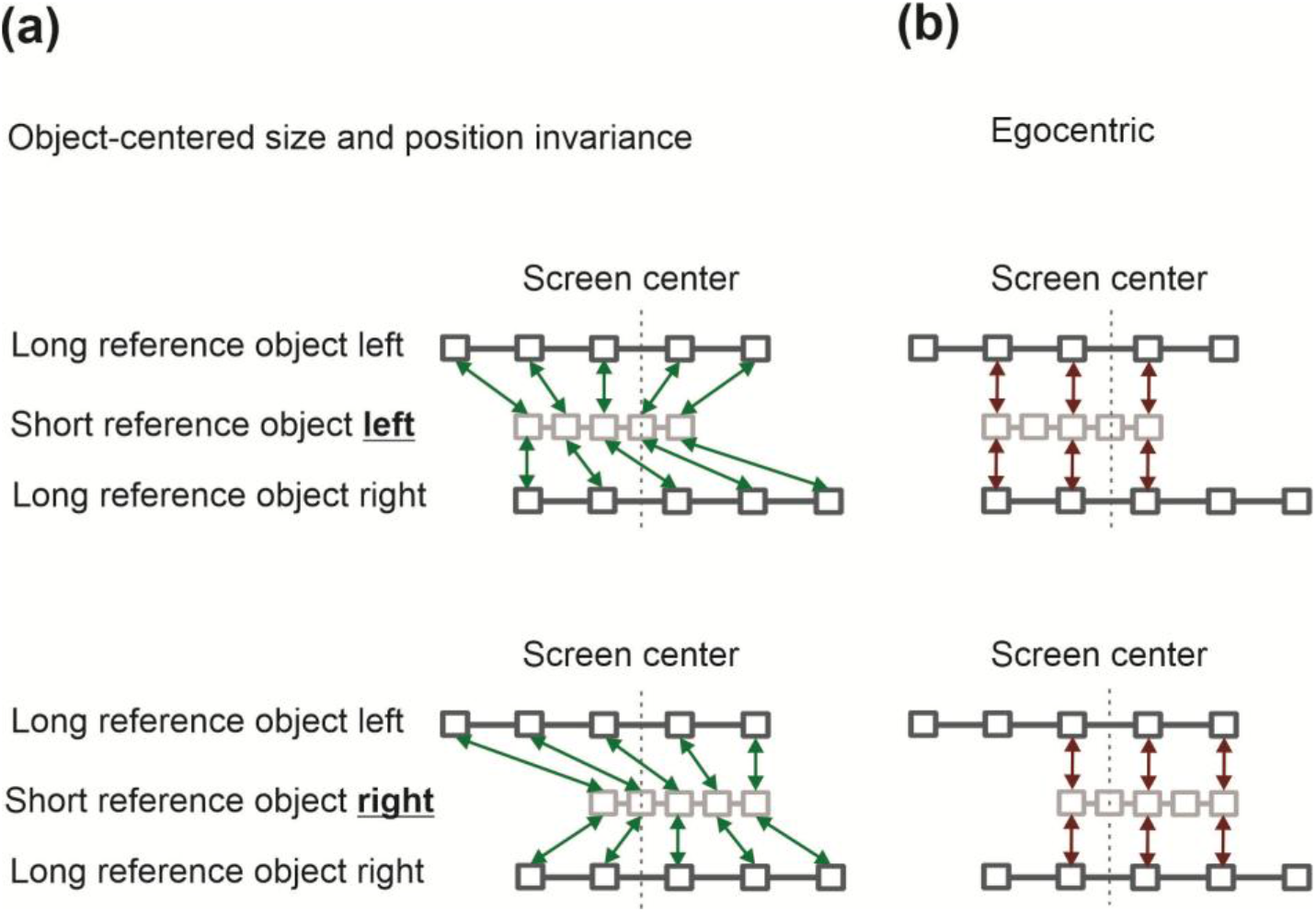
Corresponding position information for calculating Position and Size Invariance (PSI) between different task conditions. **(a) Corresponding positions for calculating alloCorr.** Invariance with respect to object position and object size predicts similar neural selectivity profiles between short-object and long-object conditions when the profile is computed relative to the within-object numerical box positions (1, 2, 3,…). To calculate alloCorr, we considered all possible correspondences (green arrows) between long and short objects when the short objects presented on the left (top panels) or right (bottom row) of the screen center (vertical dotted line). Box 1 (2, 3,…) of the object here always corresponds to box 1 (2, 3,…), irrespective of object position and size. We correlated a vector of 20 firing rates from long-object selectivity profiles with a vector of 20 firing rates from short-object profiles. Higher alloCorr results in higher PSI. **(b) Corresponding positions for calculating egoCorr.** The egocentric hypothesis predicts similar activity profiles for boxes between long and short objects which share the same position on the screen (= relative to the body). Accordingly, to calculate egoCorr, we considered every possible overlap (red arrows) between long and short objects when objects presented on the left (top row) or right (bottom row) of the screen center (vertical dotted line). We correlated a vector of 10 firing rates from long-object selectivity profiles with a vector of 10 firing rates from short-object profiles. Higher egoCorr results in lower PSI.

**Supplementary Figure 2.**
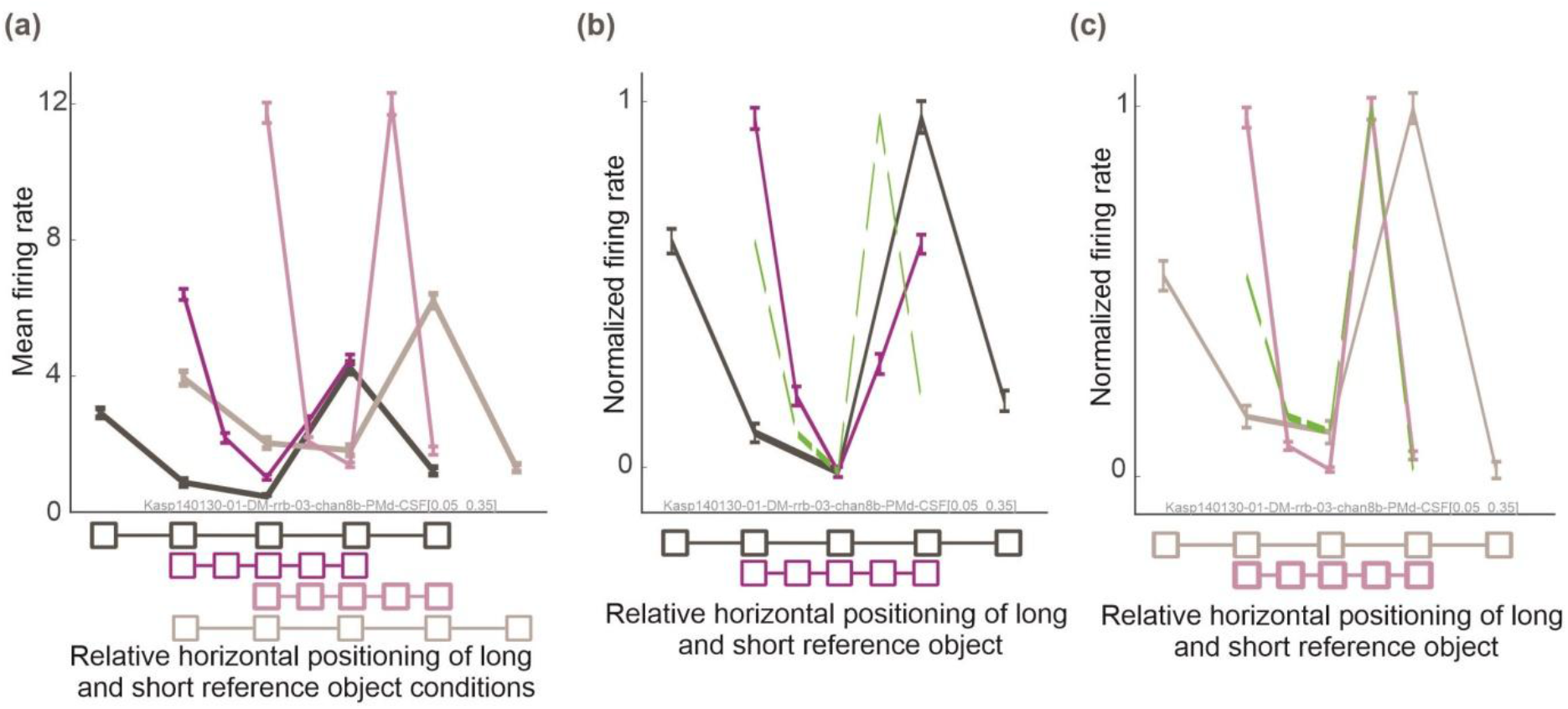
Example unit with mixed reference frame in Exp II. **(a)** Selectivity profiles of a PMd unit in early memory period, 50 – 350 ms after cue offset, for different object size and positions. **(b)** and **(c)** show normalized version of the same selectivity profiles as in **(a)**, separately for long and short objects left **(b)** and right **(c)**. The green dashed curves show the predicted selectivity profile for the short object trials based on the based on the selectivity profiles in the long object trials, assuming ideal object-centered encoding (i.e., the squeezed version of the long object selectivity profiles relative to the central box of the object). For this unit selectivity profiles in long-object-left (dark grey) and long-object-right (light grey) indicate position invariance. Also, the short-object-right (light purple) selectivity profile is quite close to a horizontally scaled version of the long-object profile, with gain modulation. In this sense, this unit meets predictions of a position- and size-invariant object-centered reference frame. However, the short-object-left profile deviates from the prediction, suggesting some form of mixed reference frame.

